# Structural equation modeling for hypertension and type 2 diabetes based on multiple SNPs and multiple phenotypes

**DOI:** 10.1101/631853

**Authors:** Saebom Jeon, Ji-yeon Shin, Jaeyong Yee, Taesung Park, Mira Park

**Affiliations:** Department of Marketing Information Consulting, Mokwon University, Daejeon, KOREA; Department of Preventive Medicine, Kyungpook National University, Daegu, KOREA; Department of Physiology and Biophysics, Eulji University, Daejeon, KOREA; Department of Statistics, Seoul National University, Seoul, KOREA; Department of Preventive Medicine, Eulji University, Daejeon, KOREA

**Author notes:** These authors contributed equally to this work (Saebom Jeon, Ji-yeon Shin). Corresponding author Email addresses (MP).

**Keywords:** genome-wide association study, hypertension, latent variable, single nucleotide polymorphisms (SNP), structural equation model, type 2 diabetes

## Abstract

Genome-wide association studies (GWAS) have been successful in identifying genetic variants associated with complex diseases. However, association analyses between genotypes and phenotypes are not straightforward due to the complex relationships between genetic and environmental factors. Moreover, multiple correlated phenotypes further complicate such analyses.

To resolve this complexity, we present an analysis using structural equation modeling (SEM). Unlike current methods that focus only on identifying direct associations between diseases and genetic variants such as single-nucleotide polymorphisms (SNPs), our method introduces the effects of intermediate phenotypes, which are related phenotypes distinct from the target, into the systematic genetic study of diseases. Moreover, we consider multiple diseases simultaneously in a single model. The procedure can be summarized in four steps: 1) selection of informative SNPs, 2) extraction of latent variables from the selected SNPs, 3) investigation of the relationships among intermediate phenotypes and diseases, and 4) construction of an SEM. As a result, a quantitative map can be drawn that simultaneously shows the relationship among multiple SNPs, phenotypes, and diseases.

In this study, we considered two correlated diseases, hypertension and type 2 diabetes (T2D), which are known to have a substantial overlap in their disease mechanism and have significant public health implications. As intermediate phenotypes for these diseases, we considered three obesity-related phenotypes—subscapular skin fold thickness, body mass index, and waist circumference—as traits representing subcutaneous adiposity, overall adiposity, and abdominal adiposity, respectively. Using GWAS data collected from the Korea Association Resource (KARE) project, we applied the proposed SEM process. Among 327,872 SNPs, 24 informative SNPs were selected in the first step (p<1.0E-05). Ten latent variables were generated in step 2. After an exploratory analysis, we established a path diagram among phenotypes and diseases in step 3. Finally, in step 4, we produced a quantitative map with paths moving from specific SNPs to hypertension through intermediate phenotypes and T2D. The resulting model had high goodness-of fit measures (*χ*^2^= 536.52, NFI=0.997, CFI=0.998).

## Introduction

Hypertension and type 2 diabetes (T2D) are two of the leading risk factors for atherosclerotic cardiovascular disease, which is a major component of the global burden of disease [1–4]. These conditions often occur together, and recent studies showed that the presence of T2D increased the risk of hypertension [5, 6]. Hypertension and T2D are thought to share common pathways such as obesity, insulin resistance, inflammation, oxidative stress, and mental stress [7]. In addition to lifestyle and environmental factors, genetic factors have also been explored to understand the mechanisms of T2D and hypertension [7, 8]. In particular, since obesity-related phenotypes are thought to be a common pathophysiological element underlying T2D and hypertension, understanding the connections among these diseases and factors related to obesity is an important aspect of the search for proper treatments of these diseases.

Genome-wide association studies (GWAS) have been successful in identifying genetic variants associated with complex diseases such as asthma, autism disorder, T2D, and hypertension [9]. A typical approach of GWAS is to report significant single-nucleotide polymorphisms (SNPs) affecting disease; in other words, such studies generally present single-SNP analyses. However, association analyses between genotypes and traits are not straightforward due to the complex relationships between genetic and environmental factors. Moreover, in the presence of multiple correlated phenotypes and/or diseases, such analyses become more complicated. Since a separate analysis of each phenotype or disease ignores dependencies among phenotypes, a multivariate approach considering a joint analysis should be considered. Various genomic studies have been conducted to understand hypertension and T2D individually [10, 11]. However, few studies have attempted to model the pathways underlying hypertension through obesity-related traits and T2D [12].

In this study, we present a structural equation modeling (SEM)-based approach to studying the associations between genetic variants and phenotypes. SEM is a multivariate statistical method that involves the estimation of parameters for a system of simultaneous equations [13]. In general, SEM is a confirmatory approach rather than an exploratory approach, in that the main focus of SEM is to verify that a model established by a researcher beforehand is supported by the data [14], although it may sometimes seem exploratory since the model can be modified to improve its goodness of fit. SEM involves representing and estimating a complex structural relationship among variables and latent constructs. It can also be used to test the hypothesized patterns of directional and non-directional relationships among observed and latent variables [15]. A useful aspect of SEM is that it can be utilized to examine relationships between latent variables, as well as the associations among latent variables and measurement variables.

Various types of models can be used in SEM, including regression, path, confirmatory factor, and growth curve models [16]. SEM has been used in various fields, including genetic analysis [17, 18]. For metabolic traits, [19] the SEM approach has been used to scan the genome for loci. Additionally, Lozano et al. [2] collected cross-sectional data and provided a path diagram from genome variants to metabolic syndrome and metabolic disease. In that study, they used SNPs that had significant associations with phenotypic traits as exogenous predictors. Meanwhile, approaches to analyzing longitudinal data with pedigrees have also been developed [20, 21]. However, few studies have applied SEM to analyze genetic information.

In this respect, we suggest a procedure for constructing an SEM to investigate the relationships of multiple SNPs and multiple intermediate phenotypes with respect to multiple diseases. Unlike current methods that focus only on identifying direct associations between diseases and SNPs, our method introduces the effects of intermediate phenotypes. We define intermediate phenotypes as disease-related phenotypes distinct from the target itself. Moreover, we simultaneously consider multiple diseases in the model. As a result, we generated a quantitative map simultaneously showing the relationships among multiple SNPs, phenotypes, and diseases.

Our detailed goals were to answer the following research questions. First, do genetic characteristics such as SNPs affect intermediate phenotypes and/or diseases? Second, is there any commonality or similarity between the SNPs that are significantly associated with each intermediate phenotype or disease? Furthermore, if there is a genetic commonality with a significant impact on a trait, can we consider their underlying component, by taking that commonality into account? How do such components affect intermediate phenotypes and disease? Third, are there any associations or causal relationships between the intermediate phenotypes and diseases under consideration? Fourth, is it possible to simultaneously consider the relationships of multiple SNPs, intermediate phenotypes, and diseases? To answer these questions, we developed an SEM-based procedure that introduced intermediate phenotypes to examine both direct and indirect relationships. We applied this approach to Korean GWAS data. We considered two diseases (hypertension and T2D) and three intermediate variables related to obesity (subscapular skin fold thickness [SUB], body mass index [BMI], and waist circumference [WC]). Using Korean GWAS data, we constructed a model with paths extending from genetic variants to hypertension through obesity-related intermediate phenotypes and T2D.

## Materials and Methods

### SEM-based modeling procedure of multiple SNPs and multiple phenotypes

In order to investigate the multiple phenotypes that reflect joint action of multiple SNPs, we developed an SEM-based modeling procedure summarized in the following four steps. Consider an n × c data matrix A with three blocks for n samples: A= [X|Y|Z]. The first block X contains data from n × p SNPs, Y is a block corresponding to n × q intermediate phenotypes, and Z is a block corresponding to n × r diseases, where c = p + q + r.

The first step was to select the preliminary informative SNPs. To avoid computational complexity and multicollinearity due to the enormous scale of the SNPs in GWAS, non-contributing SNPs to each phenotype were excluded through a single-SNP analysis using regression or logistic regression models. Since intermediate phenotypes and diseases are likely to be heterogeneous according to demographic factors such age and sex, analyses were conducted using demographic factors as covariates. Instead, it is possible to choose the most significant SNP in each linkage disequilibrium (LD) block.

The second step was to construct latent variables for the selected SNPs relevant to each intermediate phenotype and disease variable. These significant SNPs according to phenotype may have some similarities, which can be considered as common latent variables, inferred to have a similar genetic function or to be located near the same genes. For this reason, we conducted exploratory factor analysis using the informative SNPs and phenotypes. In order to construct latent variables efficiently, SNPs with a very low communality were excluded. Here, communality was considered to represent the variance of any SNP that was shared with other SNPs via common factors.

The third step was to investigate the effect of intermediate phenotypes on diseases, or the associations among all these phenotypic variables. This association analysis of multiple phenotypes provided a conceptual framework for constructing an SEM structure in the following step.

In the final step, we applied SEM based on the previously constructed latent variables, or joint SNPs, obtained from step 2. The SEM reflected the relationships between all SNPs, their joint action or latent variables, intermediate phenotypes, disease, and morbidity. A typical process for SEM was performed [13]. That is, after identification of the model, we estimated and tested the model. The model was modified if necessary. Finally, we identified the best SEM model with the highest goodness-of-fit measures. Fig 1 shows a summary of the proposed procedure.

**Fig 1.**
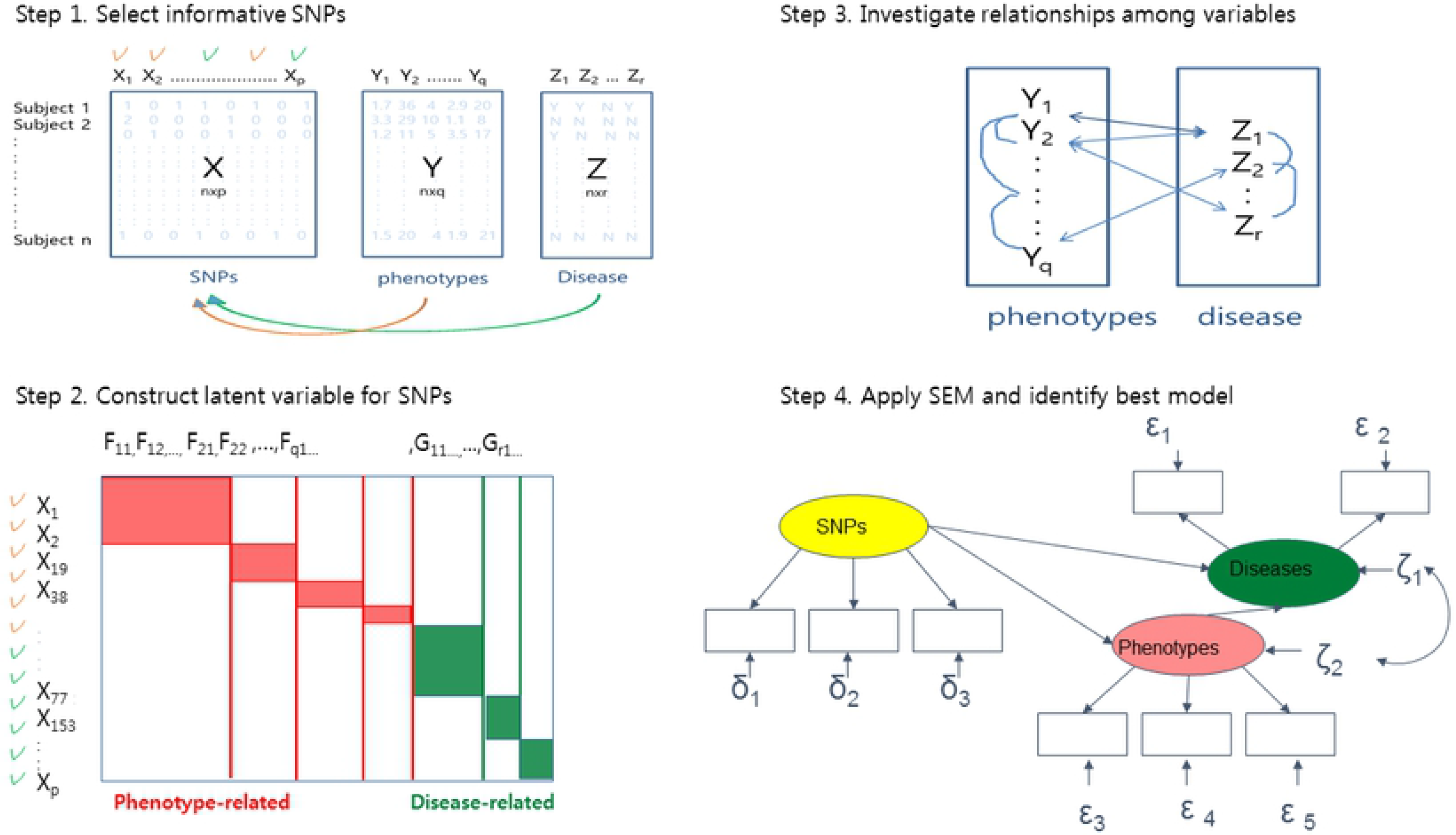
Summary of SEM-based modeling procedures for genomic data. *F*_*ij*_ represents the j^th^ factor loading for the i^th^ phenotype, *G*_*ij*_ represents the j^th^ factor loading for the i^th^ disease.

### GWAS data and phenotypic measurements

We analyzed the GWAS data set from the Korea Association Resource (KARE) project. This project was initiated in 2007 in order to undertake a large-scale GWAS. The 10,038 participants were recruited from two community-based cohorts: Ansung, representing a mainly rural community, and Ansan, representing an urban community [22]. After standard quality control procedures for the subjects and SNPs, a total of 8,842 participants and 327,872 SNPs remained. Of them, 4,183 (47.31%) were male and 4,659 (52.69%) were female, with a mean age of 52.22 years (range, 39-70 years). The KARE data include demographic characteristics such as area, sex, and age, as well as multiple phenotypes related to obesity, T2D, and hypertension that were identified based on a review of the participants’ medical records. Our research interests focused on T2D and hypertension as mediated through obesity. Table 1 presents the basic statistics for the data.

**Table 1.** Descriptive statistics of the KARE data.

| Variables | | | N (%) or mean ± SD |
| --- | --- | --- | --- |
| Demographic variables | Sex | Male | 4,183 (47.31%) |
|  |  | Female | 4,659 (52.69%) |
|  | Area | Ansung | 4,205 (47.56%) |
|  |  | Ansan | 4,637 (52.44%) |
|  | Age |  | 52.22 (8.92) |
| Intermediate phenotypes | Body mass index, kg/m<sup>2</sup> | | 24.60 ± 3.12 |
| | Subscapular skin fold thickness, mm | | 23.69 ± 10.96 |
| | Waist circumference, cm | | 82.67 ± 8.79 |
| Diseases | Hypertension | Yes | 2,393 (27.06%) |
|  |  | No | 6,449 (72.94%) |
|  | T2D | Yes | 836 (9.45%) |
|  |  | No | 8,006 (90.55%) |
| Disease component variables | Systolic blood pressure | | 117.59 ± 18.28 |
| | Diastolic blood pressure | | 75.07 ± 11.56 |
| | Fasting blood glucose | | 87.66 ± 21.88 |
| | Blood glucose after 2 hours | | 126.76 ± 51.03 |

KARE, Korea Association Resource; T2D, type 2 diabetes.

We defined T2D using fasting blood glucose (FBG) or blood glucose after 120 minutes (OGTT120), with criteria of FBG ≥126 mg/dL, OGTT120 ≥200 mg/dL, or the use of antidiabetic medication. Hypertension was defined as systolic blood pressure (SBP) ≥140 mm Hg, diastolic blood pressure (DBP) ≥90 mm Hg, or the use of antihypertensive medication. As intermediate phenotypes, we considered three traits related to obesity: BMI, WC, and SUB. More specifically, BMI reflects overall body adiposity [23], whereas WC reflects abdominal adiposity (for which visceral adipose tissue is largely responsible), and SUB reflects subcutaneous adiposity [23]. Height (cm), body weight (kg), and waist circumference (cm) were measured using standard methods in light clothes. BMI was calculated as the weight divided by the square of height (kg/m^2^). SUB was measured using a caliper at a vertical fold taken 1 inch below the lowest point of the shoulder blade (mm).

Hypertension was present in 2393 (27.06%) participants, and 836 (9.45%) of participants had T2DM. The average(± SD) values of the obesity-related variables were 24.60 kg/m^2^ (±3.12 kg/m^2^) for BMI, 23.69 mm (± 10.96 mm) for SUB, and 82.7 cm (± 8.79 cm) for WC (Table 1). We analyzed these data using the proposed procedure. From step 1 to step 3, statistical analyses were done using SAS version 9.4 (SAS Corp., Cary, NC, USA). For the SEM analysis in step 4, Lisrel version 9.1 (Scientific software international, Skokie, IL, USA) was used. The threshold for statistical significance was set at α=0.05, unless otherwise noted.

## Results

### Step 1: Associations between single SNPs and phenotypes

Among the 327,872 SNPs in the KARE dataset, we selected informative SNPs affecting intermediate phenotypes and diseases. We conducted single-SNP analyses using simple linear regression models and logistic regression models for intermediate phenotypes and diseases, respectively. Sex, age, and area were used as covariates. The SNPs were regarded as statistically significant when they showed a p-value less than 1.0E-05 in the single-SNP analysis. The obesity-related phenotypes (BMI, SUB, and WC) were associated with 19 SNPs: 7 SNPs for BMI, 5 SNPs for SUB, and 7 SNPs for WC. For the dichotomous definition of T2D, a single SNP was determined to be significant, with a p-value less than 1.0E-05. For the dichotomous definition of hypertension, 4 SNPs were identified as significant. In total, we considered 24 SNPs related with obesity, T2D, and hypertension. Table 2 presents a list of the selected SNPs.

**Table 2.** Significant SNPs for each phenotype through single-SNP analyses.

| Traits | SNP | p-value | Traits | SNP | p-value |
| --- | --- | --- | --- | --- | --- |
| BMI | rs527248 | 2.98E-06 | Waist circumference (WC) | rs4667458 | 4.72E-06 |
|  | rs17178527 | 2.02E-08 |  | rs11933222 | 9.15E-06 |
|  | rs11000212 | 1.45E-06 |  | rs17178527 | 2.71E-06 |
|  | rs7193144 | 3.30E-06 |  | rs17092358 | 6.21E-06 |
|  | rs8050136 | 2.68E-06 |  | rs2074356 | 7.95E-06 |
|  | rs9926289 | 2.45E-06 |  | rs17089409 | 7.07E-06 |
|  | rs9939609 | 1.43E-06 |  | rs17089410 | 5.67E-06 |
| Subscapular skin fold (SUB) | rs17248901 | 1.92E-07 | T2D | rs11131794 | 5.09E-06 |
|  | rs6561930 | 2.85E-07 | Hypertension | rs17249754 | 3.15E-07 |
|  | rs16951883 | 4.66E-07 |  | rs7136259 | 4.03E-07 |
|  | rs7193144 | 4.65E-06 |  | rs2254613 | 6.75E-06 |
|  | rs8050136 | 3.78E-06 |  | rs1378942 | 8.02E-06 |

SNP, single-nucleotide polymorphism; T2D, type 2 diabetes.

### Step 2: Construction of latent variables for SNPs

To identify the underlying components, we conducted exploratory factor analysis. During factor analysis, we investigated the communality between SNPs and constructed latent variables, and excluded three SNPs with very low communality (less than 0.3): rs17178527, which was related to BMI; rs16951883, which was related to SUB; and rs17092358, which was related to WC. A separate factor analysis was performed for each phenotype to obtain latent variables for SNPs. Table 3 shows the factor loadings and variance explained for each phenotype. By using factor loadings, the four SUB-related SNPs were constructed as two latent variables (hereafter, LSUB1 and LSUB2). The six BMI-related SNPs were also constructed as two latent variables (hereafter, LBMI1 and LBMI2), and the seven WC-related SNPs were composed of three latent variables, (hereafter, LWC1, LWC2, and LWC3). These latent variables may reflect the common joint action of SNPs on their respective phenotype. The latent variables for the SNPs related to hypertension (hereafter, LHYP1 and LHYP2) and diabetes (hereafter, LDIA1) were also generated in a similar manner. The SNPs classified as corresponding to the same latent variable are shown in square brackets. The two SNPs comprising LHYP2 have different signs, meaning that they were found to be related in different directions. Both LSUB1 and LBMI1 consisted of SNPs from the *FTO* gene, which is known to be associated with fat mass and obesity. The SNPs comprising LHYP1 were from *ATP2B1*, which has been reported to be a hypertension-related gene [24].

**Table 3.**
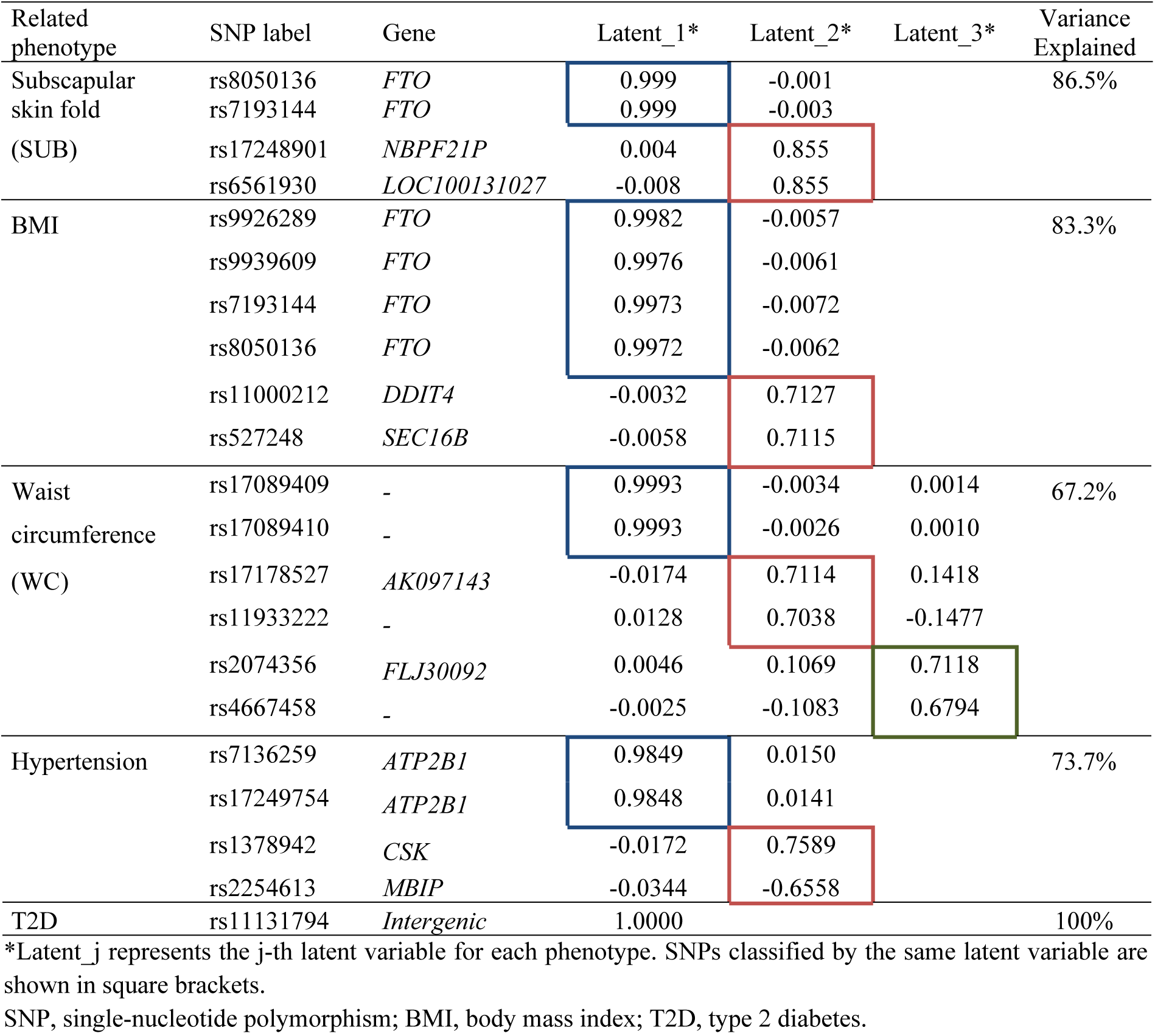
Latent variable construction of multiple SNPs for each phenotype.

### Step 3. Investigation of the relationships among variables

We investigated the association among phenotypes by adjusting for the effect of covariates including area, sex, and age. Partial correlation coefficients of the considered phenotypes are presented in Table 4. All three intermediate phenotypes were directly correlated with each other (r>0.547). For hypertension, BMI showed the largest correlation coefficient of the obesity measures (r=0.194), and WC showed the next largest correlation coefficient (r=0.180). For T2D, WC showed the largest correlation coefficient (r=0.122), implying that central body fat may be more closely associated with components of metabolic syndrome, such as T2D and hypertension.

**Table 4.** Partial correlation coefficients among obesity-related phenotypes for T2D and hypertension (n=8792)

|  | Subscapular skin fold | BMI | WC | T2D |
| --- | --- | --- | --- | --- |
| BMI | 0.634*** |  |  |  |
| WC | 0.547*** | 0.824*** |  |  |
| T2D | 0.121*** | 0.107*** | 0.122*** |  |
| Hypertension | 0.135*** | 0.194*** | 0.180*** | 0.102*** |

\*\*\* $p < 0.0001$
T2D, type 2 diabetes; BMI, body mass index; WC, waist circumference.

Before the main SEM analysis, we conducted a path analysis between the intermediate phenotypes and disease to determine the most proper model. First, we set T2D as a risk factor for hypertension according to recent studies [5, 6]. Next, we set up two major sets of paths regarding obesity indicators. The first set of paths directly linked WC to T2D or hypertension because clinical evidence suggests that the association of insulin resistance (which is thought to be the key common pathway of T2D and hypertension) with WC is stronger than the corresponding associations with BMI or SUB [22, 24]. In the second set of paths, as BMI and SUB showed considerable correlations with WC, we established distinct paths that connected BMI and SUB (1) via WC to T2D or hypertension and (2) directly, not via WC. As a result, we considered the path from obesity to T2D and hypertension that is shown in Fig 2. It displays the coefficient estimates of the path analysis, which indicate the strength of the effect of the independent variable on the dependent variable. Paths with solid lines indicate that effects were statistically significant (p<0.05), whereas two paths with dotted lines showed statistically insignificant effects.

**Fig 2.**
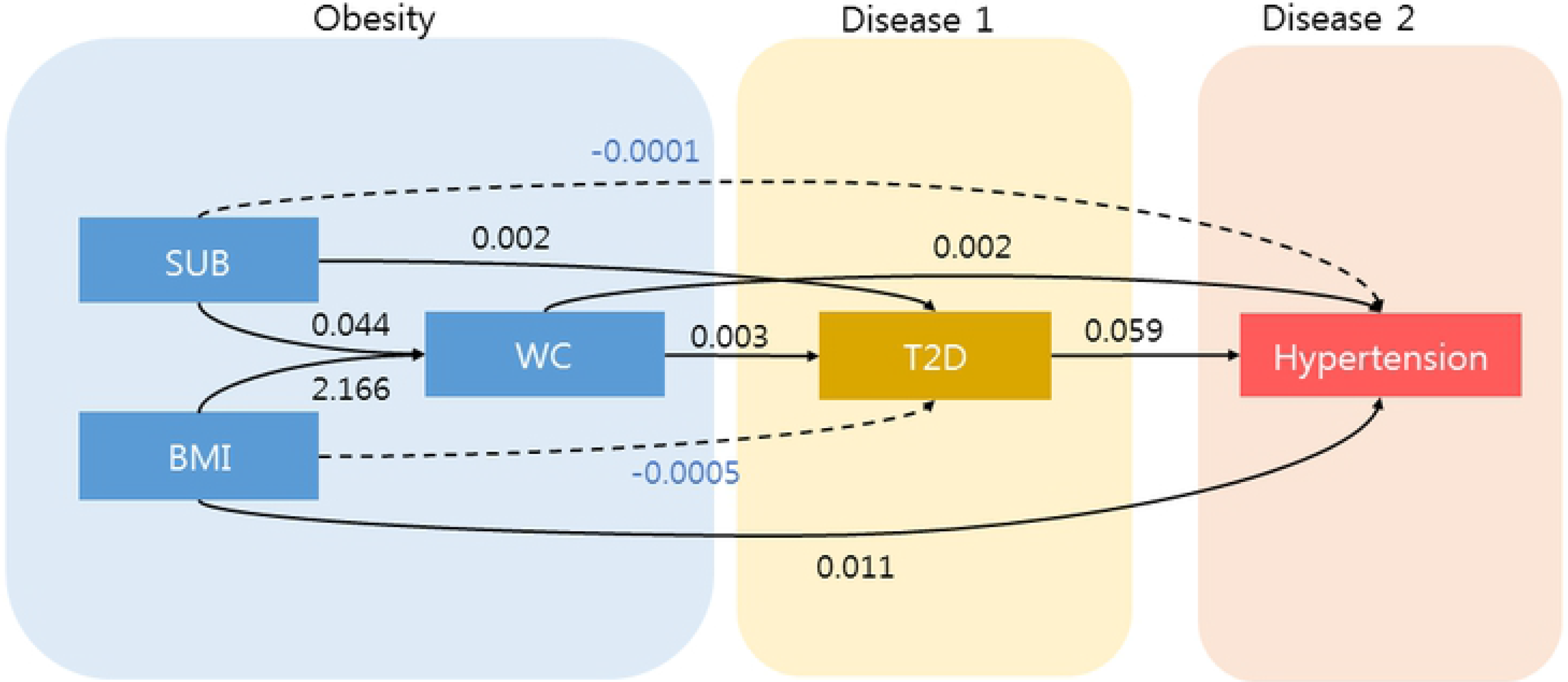
Path diagram and direct effect estimates from the path analysis of phenotypes. Solid lines indicate that effects were statistically significant (p<0.05), whereas dotted lines present statistically insignificant effects.

### Step 4: Structural equation model of multiple SNPs and multiple phenotypes

The SEM was constructed based on the joint action of the multiple SNPs and multiple intermediate phenotypes identified in the previous steps. By developing an SEM, we reflected the relationships between all SNPs, their joint action or latent variables, obesity-related phenotypes, and their possible morbidity (i.e., their effect on diseases). The model was estimated using sex, age, and area as covariates, and the fit of the hypothesized SEM was improved by allowing measurement errors to correlated for rs9926289 and rs9939609, rs7193144 and rs8050136, rs7193144 and rs7193144, rs8050136 and rs8050136. Through the modification process, we identified the best SEM model based on various goodness-of-fit measures (*χ*^2^= 536.52, NFI=0.997, CFI=0.998, GFI=0.995, AGFI=0.993, RMSEA=0.012), where NFI is the normed fit index, CFI is the comparative fit index, GFI is the goodness of fit index, AGFI is the adjusted goodness of fit index, and RMSEA is the root mean square error of approximation.

Fig 3 and Table 5 show the standardized effects of the final SEM. In the analysis of causal relationships among intermediate phenotypes and diseases, subcutaneous adiposity showed a direct, statistically significant relationship for overall adiposity (b=0.632, t=76.25), abdominal adiposity (b=0.041, t=5.23) and T2D (b=0.097, t=6.25), but statistical significance was not reached for hypertension (b=0.01, t=0.285). Overall adiposity directly affected abdominal adiposity (b=0.797, t=101.79), T2D (b=−0.024, t=−2.10), and hypertension (b=0.140, t=4.59). Abdominal adiposity (WC) directly affected T2D (b=0.097, t=4.61) and hypertension (b=0.052, t=7.44). Solid lines in Fig 3 indicate that effects were statistically significant (p<0.05), whereas dotted lines present statistically insignificant effects.

**Fig 3.**
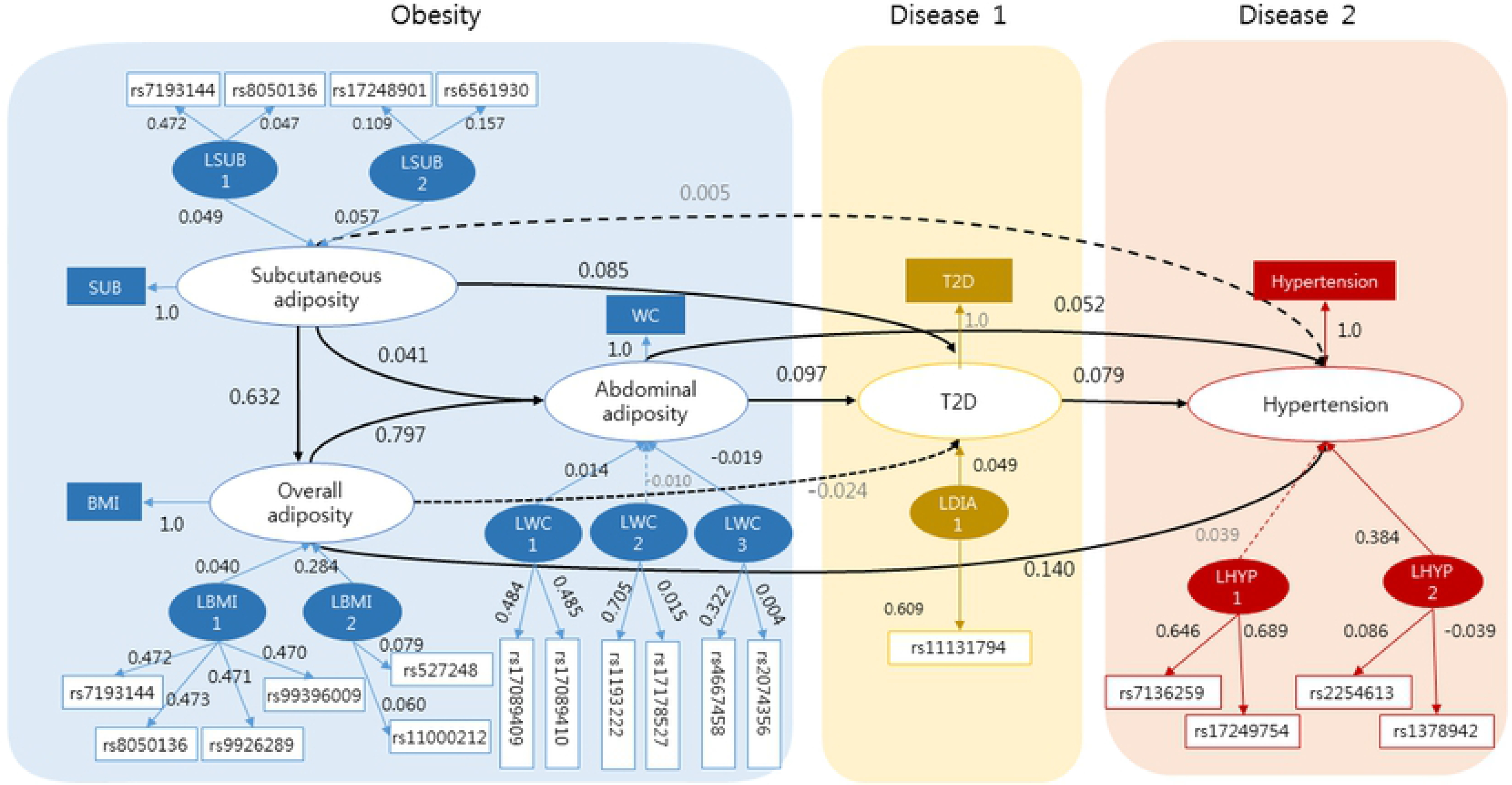
Path diagram and direct effect estimates of the final SEM. LSUBj, LWCj, LBMIj, LDIAj, and LHYP represent the j-th latent variables of the SNPs related to subscapular skin fold, waist circumference, BMI, T2D, and hypertension, respectively. Solid lines indicate that effects were statistically significant (p<0.05), whereas dotted lines present statistically insignificant effects.

**Table 5.**
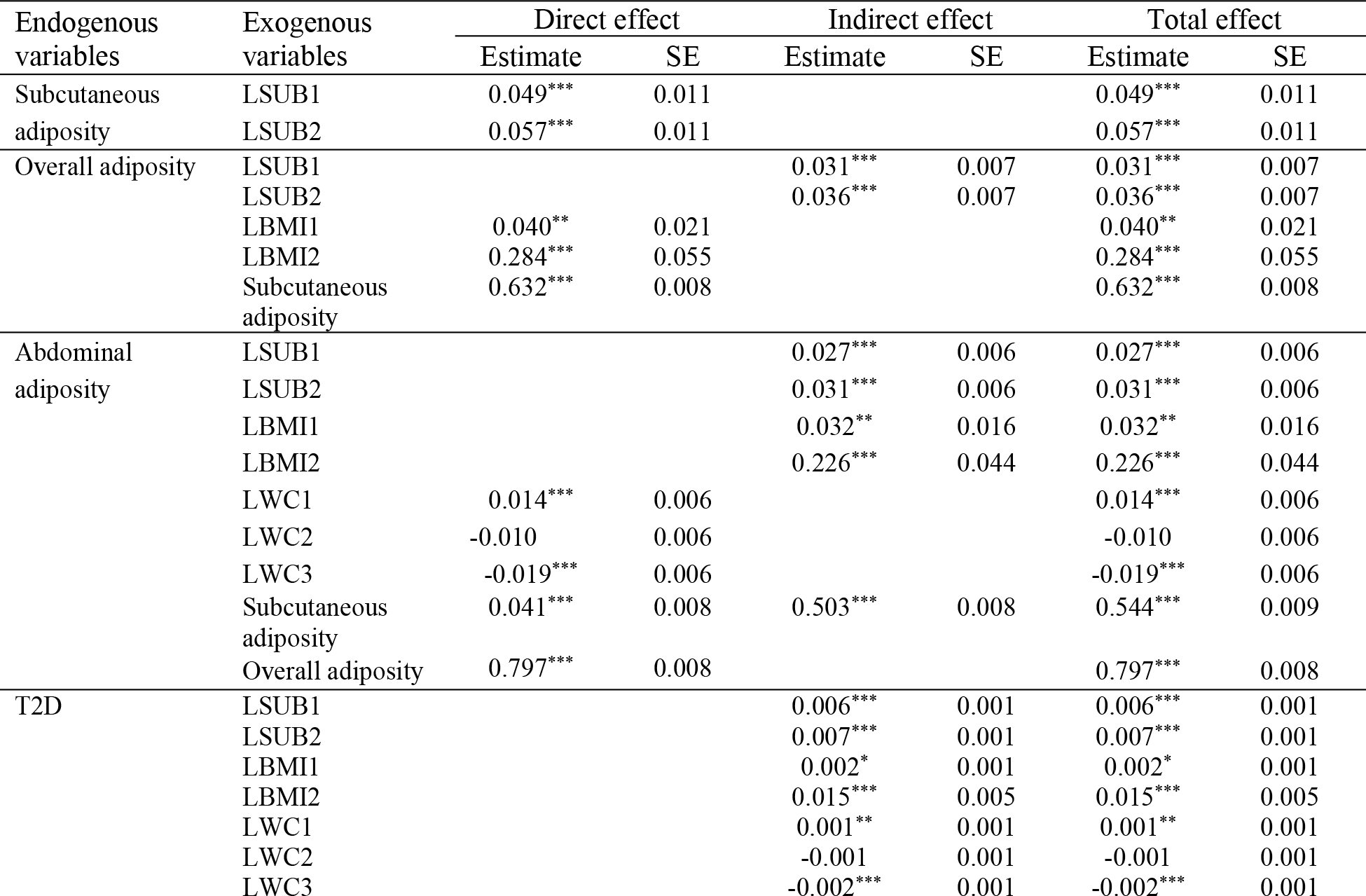
Direct, indirect, and total effects of the final SEM.

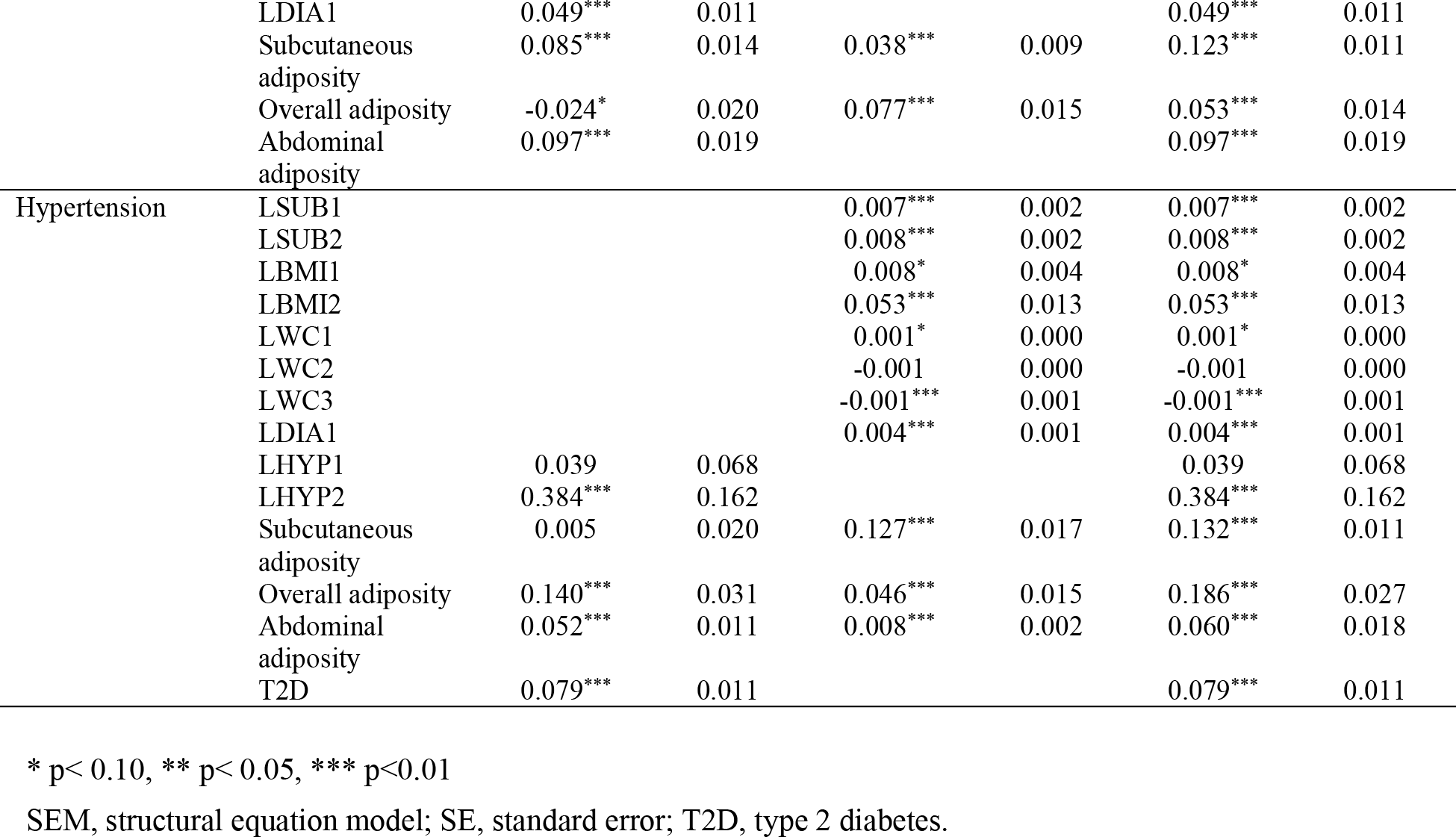

In contrast, in the analysis of genetic factors, information from 16 SNPs was used to predict T2D and information from 24 SNPs was used to predict hypertension. LSUB1 and LSUB2, which were latent variables consisting of SUB-related SNPs, showed significant direct effects on subcutaneous adiposity and also had significant indirect effects on overall adiposity, abdominal adiposity, T2D, and hypertension. Similarly, the latent variables from the SNPs related to BMI (LBMI1 and LBMI2) affected overall adiposity directly and affected abdominal adiposity indirectly. LBMI1 also significantly affected T2D and hypertension indirectly. Of the latent variables from the SNPs related to WC, LWC1 and LWC3 showed significant relationships with abdominal adiposity and T2D, but LWC2 did not. Of the WC-related latent variables, only LWC3 showed significance for hypertension. One of the latent variables from the hypertension-related SNPs (LHYP2) was significantly associated with hypertension, but the other (LHYP1) was not. However, although LHYP1 consisted of SNPs that had significant individual effects on hypertension in the single-SNP analysis, the latent variables were no longer significant when the various factors were considered together.

Integrating these observations, T2D was found to be significantly affected by subcutaneous adiposity, overall adiposity, and abdominal adiposity, and all the latent variables from the SNPs except LWC2. The largest direct effect on T2D came from abdominal adiposity (b=0.097, t=5.21). However, including indirect effects, subcutaneous adiposity had the largest total effect (b=0.123, t=11.60). Meanwhile, hypertension was directly and indirectly associated with overall adiposity, abdominal adiposity, and T2D. Subcutaneous adiposity significantly affected hypertension indirectly, but not directly. The largest effect on hypertension was obtained for LHYP2 (b=0.384, t=2.37), and overall adiposity showed the second largest direct effect (b=0.186, t=6.88). The results were similar when only direct effects were considered.

## Discussion

Hypertension and T2D are important public health concerns, as the prevalence of each is increasing worldwide [25, 26]. The coexistence of hypertension and T2D dramatically increases the risk (2- to 4-fold) of cardiovascular disease and all-cause death [4]. Although studies have investigated the effects of obesity-related factors on T2D and hypertension separately, few studies have investigated the pathways underlying hypertension through obesity-related traits. In this study, we aimed to improve our understanding of the pathways underlying hypertension and T2D driven by genetic variants and obesity-related traits by conducting a multivariate analysis. In order to achieve these goals, we developed an analytical process consisting of four steps that yielded successful results. In step 1, we investigated GWAS variants that affected intermediate phenotypes and disease, the first research goal. In step 2, we found communalities or similarities among SNPs for each phenotype, the second research goal. Step 3, the path analysis of multiple intermediate phenotypes and diseases, suggested plausible associations among traits, the third research goal. Step 4 enabled us to achieve our final research goal through an SEM analysis of the associations among multiple SNPs, multiple phenotypes, and multiple diseases. As the final result, we developed a quantitative map simultaneously showing the relationships among GWAS variants, intermediate phenotypes, T2D, and hypertension. This analysis provides insights into the mechanisms underlying T2D and hypertension. Our findings highlight the importance of subcutaneous adiposity and abdominal adiposity, as well as latent variables from SNPs, as driving elements of T2D in the Korean population. The impacts of latent variables of the SNPs, overall adiposity, abdominal adiposity, and T2D on hypertension were also confirmed. The resulting model had high goodness-of-fit measures.

## Acknowledgements

This research was supported by the Bio & Medical Technology Development Program of the NRF funded by the Korean government, MSIP (No. 2016M3A9B694241). This research was supported by Basic Science Research Program through the National Research Foundation of Korea (NRF) funded by the Ministry of Science, ICT & Future Planning (NRF-2017R1A2B4011504).

